# A deconvolution solution to aliasing confusion in apparently undersampled oblique plane microscopy

**DOI:** 10.1101/2025.04.30.651458

**Authors:** Jacob R. Lamb, Conor McFadden, Reto Fiolka, James D. Manton

## Abstract

In oblique plane microscopy, a remote refocussing system is used to create an aberration-free 3D image of the object, from which an inclined plane is sampled using a tilted imaging system. As a result, unlike in conventional microscopes, the point spread function (PSF) and optical transfer function (OTF) are tilted with respect to the optical axis of the imaging system. This suggests that a small sample-scanning step size is required to satisfy the Nyquist–Shannon sampling criterion. Recently, we have shown that a careful choice of apparent undersampling leads to aliased OTF copies that can be separated and stitched together in Fourier space to form a properly sampled OTF. In our demonstrations, this led to acquisition speed gains of between two- and four-fold. Here, we introduce a deconvolution-based method to reconstruct properly sampled volumes from apparently undersampled datasets that outperforms our previous Fourier-stitching approach.

---

Oblique plane microscopy (OPM) has established itself as a powerful light sheet fluorescence microscopy (LSFM) method with the significant advantage of conventional sample mounting [1]. In addition, the possibility of acquiring volumetric data by scanning the light sheet and image plane through the sample, rather than scanning the sample through the light sheet, can greatly accelerate volumetric imaging rates without perturbing the specimen [2–4]. However, unlike in conventional LSFM systems, the principal axes (*x, y, z*) of an OPM point spread function (PSF) are not all aligned with the sampling axes (*s*_*x*_, *s*_*y*_, *s*_*z*_).

This is because the light sheet is launched into the sample at an oblique angle and so the image plane must also be tilted at the same angle. As such, both *s*_*x*_ and *s*_*z*_ have components in the *x* and *z* directions. This has previously led to the idea that fine step sizes are required to obtain properly sampled volumetric data, as lateral and axial resolution are related via this tilted plane. Recently, we showed that this was not the case, and that by carefully exploiting the aliasing in apparently undersampled datasets a fully sampled volume could be straightforwardly reconstructed when using larger step sizes [5]. In our demonstrations, this led to acquisition speed gains of between two- and four-fold.

In the previously introduced Fourier-stitching approach, the undersampled data is 3D Fourier transformed, with *N* − 1 copies of the Fourier spectrum concatenated along 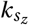. This forms a full OTF flanked by subsidiary aliased copies, which can be removed by cropping in Fourier space (implemented as convolving the data with a real-space sinc function). The fully-sampled volume can then be recovered by inverse Fourier transforming the cropped spectrum.

This method is quick but, as the spectrum is cropped, some signal is lost. Furthermore, for any real, measured volume we expect no counts to be negative and for the predominant noise statistics to be that of Poisson (shot) noise (in the common case that read noise is negligible). However, in the Fourier-stitched reconstructed volume, counts may be negative and will not obey Poisson statistics. This may compromise further downstream processing of the data, such as denoising and deconvolution. As such, it would be desirable to develop a processing approach that operates on the same data but obviates these issues.

Here, we use a Richardson–Lucy (RL) deconvolution approach as an alternative that ameliorates these problems. In the RL algorithm, the estimate of the object, *u*, is updated using the iterative rule *u*_*t+*1_ *= u*_*t*_ · H^T^(*d* /H*u*), where *d* are the measured data, H is the image formation operator (e.g. blurring with the PSF), H^T^ is the transpose (dual) of the image formation operator (e.g. blurring with the flipped PSF), and both multiplication and division operate element-wise. For our approach, we set H to be the action of blurring with the PSF and ‘throwing away’ the missing slices in the undersampled data. For the action of H^T^, we first add in empty ‘missing’ slices and then blur with the flipped PSF. After the algorithm has terminated, we are left with a deconvolved reconstruction that is fully-sampled — the fully-sampled image data can be obtained by blurring with the PSF and not throwing away any slices. Throughout, it is crucial that the PSF has the same sampling as that desired in the final reconstructed volume, otherwise an erroneous result will be produced.

To illustrate our approach, we begin with synthetic 2D data. A ground truth object (Figure 1a) is blurred with a tilted PSF (as in OPM) to produce a fully-sampled blurred image (Figure 1b). With 8× undersampling, seven slices out of eight are empty, leading to aliased copies of the blurred spectrum (Figure 1c). In reality, the undersampled data does not contain empty slices, as these are merely not recorded, but we display the full volume with missing slices for clarity. The corresponding Fourier spectrum for the undersampled dataset is contained within the cyan line. After running RL deconvolution, we obtain a reconstruction of the ground truth object (Figure 1d) where the algorithm has attempted to undo the effects of both blurring and undersampling. To obtain a fully-sampled volume, without aliasing artefacts, this reconstruction is reblurred with the fully-sampled PSF to obtain a fully sampled image volume (Figure 1e). By this process, the RL deconvolution has led to an accurate regeneration of the fully sampled, aliasing-free data from the apparently undersampled dataset.

**Figure 1:**
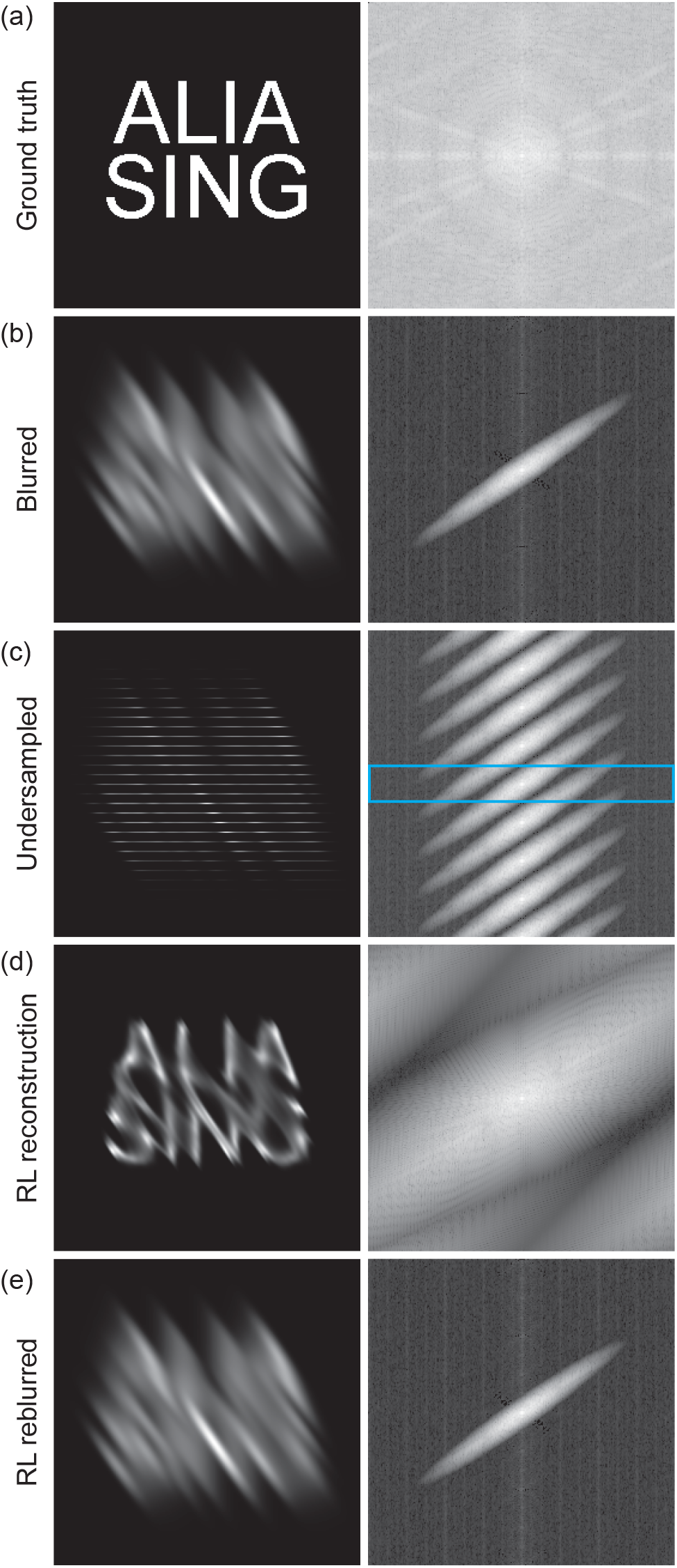
Principle of deconvolution solution to aliasing confusion. (a) Ground truth object and Fourier spectrum. (b) Blurred ground truth object and Fourier spectrum. (c) 8× undersampled blurred data and Fourier spectrum. Cyan line denotes spectrum of undersampled data without missing slices. (d) RL reconstruction of ground truth object and Fourier spectrum. (e) Reblurred RL reconstruction using full (i.e. not undersampled) PSF and Fourier spectrum. Note that aliased copies of spectrum in (c) are removed and that the image appears identical to the blurred, fully sampled image.

RL deconvolution has been noted to often produce artefacts in the reconstructed ground truth object due to the presence of noise in the blurred image, but this does not affect the quality of the reblurred reconstruction. Indeed, these artefacts arise from an overfitting of the ground truth reconstruction to the pixel-level variations in the blurred image arising from the noise — an over-deconvolved reconstruction actually leads to a better match to the original noisy blurred data when reblurred. Furthermore, when deconvolution is terminated before the result is overfit to the noise, our method acts a denoiser with the reblurred reconstruction having a less noisy appearence than the original raw data. As the deconvolution process maintains total signal levels, an appropriately noisy reconstructed dataset can be generated by adding Poisson noise to the reblurred deconvolution result (assuming that an *n*-fold undersampled dataset contains *n*-fold fewer counts than a fully-sampled dataset)

This process is first demonstrated on a high-resolution OPM system with clathrin in U2OS cells as a target, imaged live at 37 °C. Data were acquired on a home-built highresolution OPM system, as used in [5]. The primary objective is a Nikon 40× 1.25 NA silicone oil lens with a 45° light-sheet tilt.

Figure 2a shows a maximum intensity projection (MIP) of a fully sampled dataset, acquired with a lateral pixel pitch of 147 nm and stage step of 210 nm, giving a post-deskew *z*step size of 148 nm (i.e. effectively isotropic voxels). The volume has been deskewed and rotated into a coverslip-centric coordinate system, such that the MIP along *z* projects into the surface of the coverslip. A merge between the fully sampled dataset in green and the same in magenta is shown beneath the diagonal slash (producing white in overlap), with a Fourier spectrum below that. In Figure 2b, a 4× undersampled dataset (producing by throwing away three slices out of four from the fully sampled dataset) has been upsampled using a bicubic interpolation, and displayed in green in the merge, with the fully sampled dataset in magenta. As evident from the green hue to the merge image, diffraction-limited objects are represented with an inferior resolution. In contrast, the RL reconstructed dataset in Figure 2c appears as a perfect match to the fully sampled dataset.

**Figure 2:**
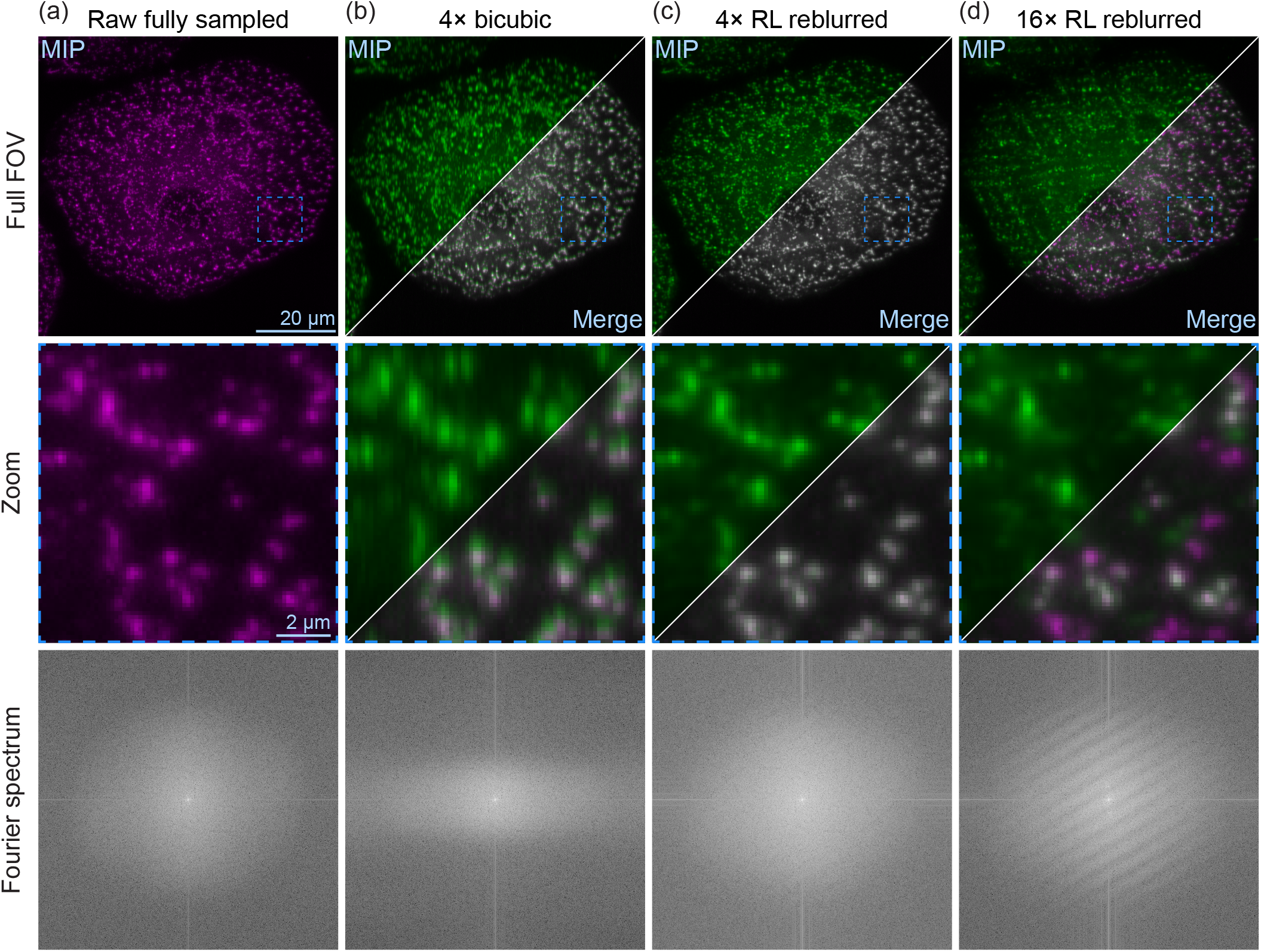
Comparison of deconvolution solution to bicubic interpolation and full sampling. Each column first shows the dataset in green, with a green-magenta merge with the fully sampled dataset shown beneath the diagonal slash, then a zoom of the cyan boxed region, and the respective Fourier spectrum at the bottom. (a) Fully sampled volume (i.e. 210 nm stage step). (b) 4× undersampled dataset (i.e. 840 nm stage step), upsampled using a bicubic interpolation. Note that the diffraction-limited clathrin clusters now appear more blurred than in the fully sampled dataset. (c) 4× undersampled dataset reconstructed using the deconvolution approach. Note the apparently perfect agreement with the fully sampled dataset. (d) 16× undersampled dataset reconstructed using the deconvolution approach. Note that there are clear artefacts from missing slices that the algorithm cannot recover.

While RL deconvolution, due to its iterative nature and positivity constraint, is known to generate spatial frequencies beyond the imaging system bandlimit, the dataset shown in Figure 2d shows that this does not lead to issues when dealing with truly undersampled data. Here, a 16× undersampled dataset (i.e. with a stage step of 3360 nm) is reconstructed using the deconvolution approach. Clear banding artefacts, in both the real space image and Fourier spectrum, can be seen, where the algorithm has accurately not been able to reconstruct data for these missing slices.

Next, we compared the deconvolution approach to the previous Fourier filtering approach using nuclei in a brain organoid. For tissue-scale imaging, data were acquired on a home-built mesoscopic OPM system, described by Lamb et al. [6]. Briefly, the primary objective is an Olympus 4× 0.16 NA lens and the system has an overall magnification of 1.347× from sample to remote space, optimised for imaging watery samples. With a light sheet NA of 0.02 and a tilt of 45°, the system achieves a full-width half-maximum resolution on 200 nm FluoSphere Yellow-Green beads of 2.05 ± 0.06 µm × 2.42±0.02 µm×22.31±0.35 µm (*n =* 12 beads). A 2 µm sample-stage shift was used for all datasets and PSF measurements, and the camera pixel pitch in the sample is 1.040 µm. A 4× UExM expanded [7] cerebral organoid, labelled with Hoescht 33342, was used as the sample.

Figure 3a shows a slice from a fully sampled dataset, with zoom beneath. Figure 3b shows a reconstruction from a 5× undersampled dataset generated using the Fourier filtering approach. While the overall morphology is correct, the image lacks Poisson noise character and there are some banding artefacts near the cyan arrows. In contrast, the RL deconvolution reconstruction shown in Figure 3c shows none of these artefacts. However, the Poisson noise character is not present, as the reconstruction has acted as an effective denoiser. If a true facsimile of the fully sampled dataset is required, appropriate Poisson noise can be added, producing the result shown in Figure 3d. Figure 3e shows the deconvolved estimate of the ground truth object produced by the deconvolution approach, which also exhibits no artefacts from the undersampling. Figure 3f demonstrates that the artefacts in the Fourier filtering reconstruction are related to the sinc kernel used to filter out aliased copies of the sample spectrum, and that no artefacts or aliased copies are present in the Fourier spectrum for the deconvolution reconstruction. Comparing reconstructions to the fully sampled dataset, we obtain root mean square errors of 23.6 for Fourier filtering, 13.8 for the RL reconstruction and 16.8 with added Poisson noise.

**Figure 3:**
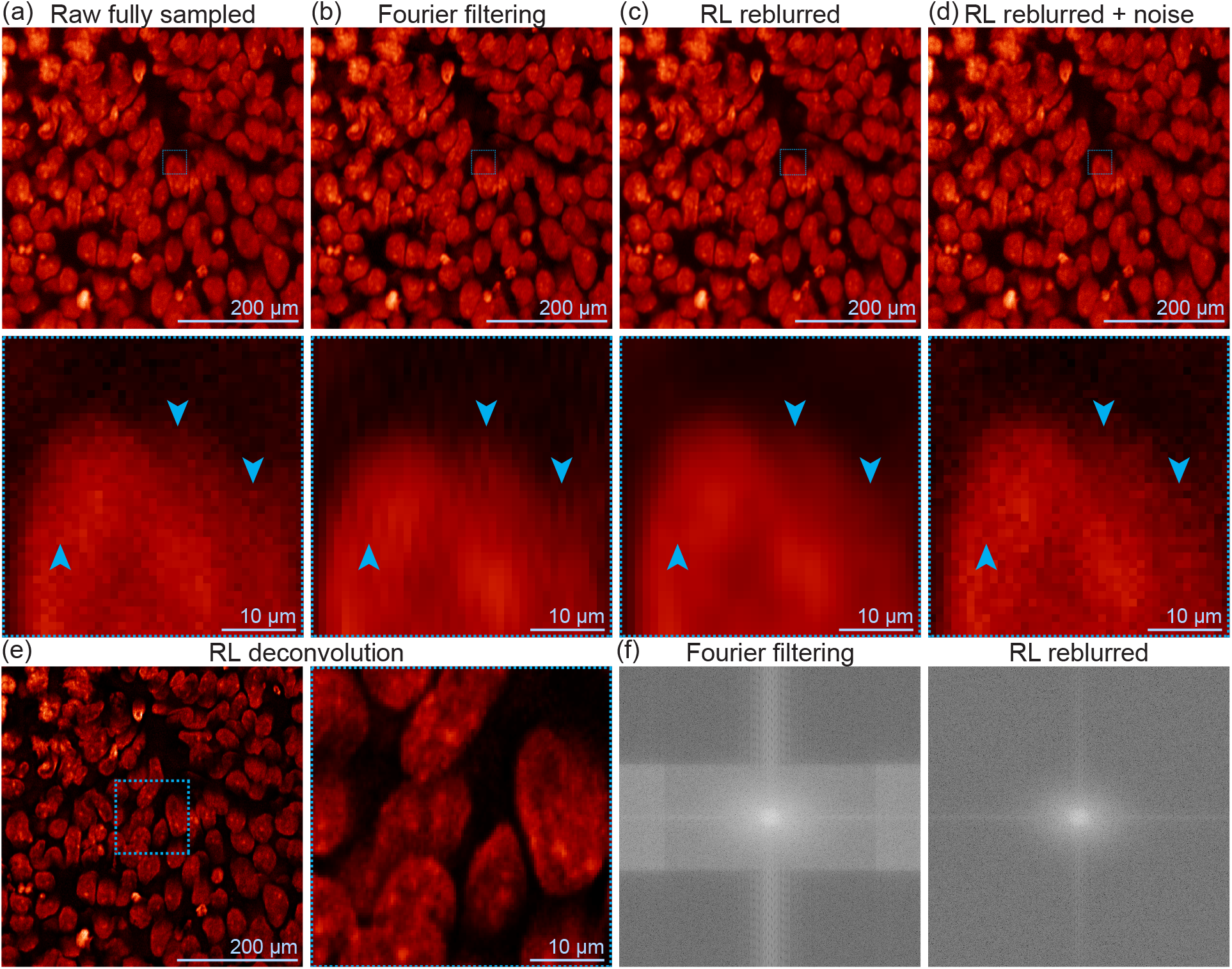
Comparison of deconvolution solution to Fourier filtering and full sampling. Each column in (a)–(d) shows a slice from the respective dataset, with a zoom of the central area shown below. Cyan arrows point to a subset of the locations of artefacts in the Fourier filtering dataset. (a) Raw, fully sampled volume (i.e. 2 µm stage step). (b) Reconstruction from 5× undersampled dataset (i.e. 10 µm stage step) using the Fourier filtering approach previously introduced by McFadden et al. [5]. Note the banding artefacts in the regions near the cyan arrows and the lack of Poisson noise character. (c) Reconstruction from 5× undersampled dataset using the deconvolution approach. Note the lack of banding artefacts in the regions near the cyan arrows and the denoised character of the image. (d) Reconstruction from (c), but with appropriate Poisson noise added. (e) Deconvolved ground truth estimate produced when generating (c). (f) Fourier spectra for the Fourier filtering and deconvolution approach datasets. Note the unusual appearance of the former due to the sinc kernel used to remove aliased copies of the true spectrum.

In summary, we have demonstrated that a simple application of RL deconvolution, followed by reblurring, can be used to recreate fully sampled OPM volumes from apparently undersampled datasets, thereby allowing for an increased acquisition rate and lower photodamage. While this iterative solution requires more computational effort than our previously introduced Fourier filtering approach, volumes produced using the new method benefit from a lack of artefacts and the side-effect that the method acts as a denoiser. Importantly, the method is quantitative and so appropriate Poisson noise can trivially be added back to the volume if desired. Furthermore, the approach necessarily produces a deconvolved ground truth reconstruction as well, as often desired for light sheet microscopy experiments to enhance contrast and apparent resolution.

## Acknowledgements

JRL and JDM thank Miguel Mestre for providing the cerebral organoid sample used in this work. This work was funded through a Royal Society University Research Fellowship to JDM (URF\R1\221086 & RF\ERE\221078).

